# FuncSeek: Multi-PLM contrastive learning for protein functional similarity search

**DOI:** 10.64898/2026.08.07.738656

**Authors:** Leendert J. Cloete, Hugh G. Patterton

## Abstract

Below 30% pairwise sequence identity, alignment-based methods struggle to reliably distinguish true homologs from chance, and enzyme function prediction degrades accordingly: on proteins in this regime, even advanced methods (CLEAN) achieves only 55.1% accuracy at full EC specificity on the CARE benchmark. To this end Protein Language Models (PLMs) have gained favor as alternatives. However, PLMs often encode only a subset of the biology, whereas the understanding of enzyme function requires among other things a combination of sequence, structure and functional-context simultaneously. In this work, we describe FuncSeek, a contrastive learning model which utilizes three diverse, complementary PLMs: ESM2 (to model evolutionary co-variation), ProstT5 (for bilingual sequence and structure embeddings), and ProteinBERT (for functional semantic similarities). Using SwissProt data, these 2816-D embeddings are labelled with Enzyme Commission numbers (EC) and are trained through a supervised contrastive head into a 256-D space. FuncSeek attains 64.6% nearest-neighbour EC4 accuracy on the CARE out-of-distribution benchmark set (*ood30*; proteins below 30% identity to training set), outperforming CLEAN (55.1%) and Diamond BLASTp (51.4%), and obtains 93.7% nearest-neighbour EC4 accuracy on the promiscuous, multi-functional enzymes benchmark (CLEAN, 69.4%). We also show that the learned representations transfer without retraining to the TrEMBL database, achieving 97.3% nearest-neighbour EC4 accuracy on an 8,031 BRENDA-validated enzyme set, never seen during training. Because only projected embeddings are stored in the target index, and function is inferred from an annotated reference set, we propose this paradigm for rapidly searching extremely large metagenomic databases, bypassing costly sequence alignment and annotation pipelines.

**Author Summary:** Enzymes are the molecular machines that carry out the chemistry of life, and knowing which reaction an enzyme performs is essential for medicine, biotechnology, and understanding how organisms work. Yet new protein sequences are being discovered far faster than we can study them in the laboratory, and the most common computational shortcut, comparing a new sequence to well-studied ones, breaks down when the sequences are only distantly related. We asked whether recent artificial intelligence models that learn the “language” of proteins could close this gap. Rather than rely on a single model, we combined three that each capture a different aspect of protein biology: evolutionary patterns, three-dimensional shape, and functional context. We then trained a system, which we call FuncSeek, to arrange enzymes so that those performing the same reaction sit close together. FuncSeek predicted enzyme function more accurately than the leading existing tools, especially for distantly related and multi-functional enzymes, and this accuracy carried over to proteins it had never seen. Because it represents each protein as a compact numerical fingerprint, it can search enormous, unexplored collections of sequences quickly.

## Introduction

The 2022_03 release of UniProt had in excess of 227 million protein sequences, with only 226,101 of these being manually reviewed. Of the remaining unreviewed entries (TrEMBL), rule-based annotation systems like UniRule and the Association-Rule-Based Annotator covered only 53.4% of these, leaving ∼55 million sequences still without any functional annotation beyond ‘Uncharacterized protein’ [1]. By the 2024_04 release, the number of entries had grown to a total of 246 million sequences and 231,709 curated entries, with ProtNLM, a large language model, renaming more than 28 million of these previously uncharacterized proteins [2]. However, the overwhelming majority of protein sequences still have no functional annotation, either manually assigned or predicted. The disparity between structural and functional coverage further delineates the problem clearly. Recently, more than 85% of UniProt entries had a predicted 3D structure derived from AlphaFold [1, 3], yet we do not know the function of the majority of these proteins.

For enzymes, this problem is even more evident. Schnoes et al. [4] found that misannotation of molecular function in enzyme superfamilies averaged anything from 5% to 63% across the six superfamilies studied (GenBank NR, TrEMBL, and KEGG), with 10 of the 37 families examined exceeding 80% misannotation in at least one database.

Despite advances in the field of functional annotation, BLASTp still remains the tool most frequently used to assign functions. It is based on the premise that similarity between two sequences indicates functional commonality [5]. But below 30% pairwise sequence identity, this premise often no longer holds, and alignment based annotation starts to struggle. Within this “twilight zone”, homology cannot be distinguished from random matching [6] and neighbour annotation transfer begins to fail. Concurrently, databases are also expanding far more rapidly than experimental verification can match [4]. As many as 65-90% of sequences in metagenomic and metatranscriptomic datasets bear no detectable similarity to any characterised organism, making functional annotation difficult [7–9].

MMseqs2 was designed to break through this computational bottleneck, achieving over 400 times the speed of PSI-BLAST, while exceeding it in iterative profile search sensitivity [7]. However, even alignment-based annotation methods cannot assign function when no annotated homolog falls within the allowed difference, regardless of its speed or sensitivity. Thus, this increasing discrepancy between the pace of sequencing and biological characterisation required the development of approaches that retrieve functional information from regions of sequence space that are inaccessible to alignment approaches.

Protein language models (PLMs) offers an alternative. ESM-1b, a 650 million parameters Transformer, trained on UniRef50 sequences via masked language modelling, demonstrated that unsupervised pre-training alone can generate biologically meaningful organization in the learned representation space. Orthologous proteins cluster together, secondary structure and tertiary contacts can be linearly interpreted, and vector nearest-neighbors searches retrieve distantly related proteins at fold-level similarity, approaching hidden Markov model based methods like HHblits [10]. Similarly, since protein structure and function constrain which mutations survive natural selection, language models trained on natural sequences learn to recover these evolutionary constraints, thereby extracting features that are informative of biological properties [10, 11]. The successful scaling of its successor (ESM-2) to 15 billion parameters enabled the development of ESMFold, which demonstrated that single sequence protein structure prediction with atomic-level accuracy and a mean TM-score of 0.83 on CAMEO was possible, without taking multiple sequence alignments as input [11]. Thus, PLMs have established themselves as a powerful general feature extractor for various biological tasks.

PLMSearch used this ability to remotely infer homology at scale: coupling ESM-1b embeddings with a structure-similarity predictor, it was able to reach sensitivity >3x that of MMseqs2 [12]. CLEAN also followed a similar approach, but on the more difficult problem of predicting enzyme function, in which two proteins may have similar structure, yet catalyse entirely different reactions. Yu et al. [5] trained a supervised contrastive head on ESM-1b representations, in order to project sequences into an embedding space in which Euclidean distance reflects functional similarity as defined by Enzyme Commission (EC) number. This contrastive approach circumvents the class imbalance problem that cripples multilabel predictors on infrequently encountered EC numbers: rather than trying to predict thousands of classes, the model simply learns a distance function. Using this approach, CLEAN reached an F1 of 0.499 on the New-392 benchmark (novel sequences released after CLEAN’s training cutoff) and 0.495 on *Price* (a curated set of previously misclassified enzymes), the latter representing a 3.0-fold improvement over ProteInfer and an almost 5.8-fold improvement over DeepEC [5].

PLMs, however, have a key limitation: the pretraining task determines which information is implicitly stored in the encoder’s embeddings, and no single pretraining task captures all aspects of protein biology. Yet, while masked language modelling captures co-evolutionary signals, enabling these models to recover residue-residue contacts, secondary- and tertiary structure without explicit supervision [11], sequence statistics alone leave much functional information inaccessible, even for models as large as 15 billion parameters (ESM2). This is evident in the fact that ESMFold achieved a TM score of 0.68 on CASP14 while AlphaFold2’s multiple sequence alignment based method scored 0.85 [11]. Shaw et al. [13] provided evidence for this limitation by reporting that embedding similarity and sequence similarity yielded disjoint sets of errors for distinguishing functions of paralogs, and concluded that sequence similarity captures evolutionary relatedness while only partially reflecting functional similarity, whereas embedding similarity captures shared sequence patterns while remaining largely blind to evolutionary history. The two signals are thus complementary rather than redundant. The CARE benchmark [14] quantifies the cost of relying on any single modality: CLEAN reached only 55.1% EC4 accuracy on remote homologs (sequence with <30% identity) and 31.8% near activity cliffs, where the structure-based Foldseek scored 41.2% [14].

Therefore, given that most of what sequence pre-training leaves inaccessible is likely structural, and since AlphaFold2 can now create high-quality structures at scale, there is an argument for incorporating a structural signal directly into the model rather than having it learn it from sequences alone. ProstT5 [15] implemented this, by fine-tuning the 3B-parameter ProtT5 to become a bilingual encoder-decoder language model, which can translate between sequence and 3Di structure (a structural alphabet for residues which describes a residue’s geometric state with respect to its nearest neighbour using 20 states [16]). By directly predicting 3Di states from sequences, it can achieve near experimental structure sensitivity on distant homology detection without 3D coordinates at inference. On the task of CATH classification, ProstT5 boosted mean accuracy from 62.0% to 73.0% using their sequence-derived embeddings, and 77.0% when using experimental 3Di structures, outperforming ESM-1b by 11 and 15 percentage points, respectively [15]. The authors also showed that sequence and structure derived representations offer partially orthogonal information, and that integrating sequence and predicted 3Di embeddings increased performance at most levels of the CATH hierarchy.

However, adding structure comes at a cost. Some functional tasks slightly worsened when the model was fine-tuned, in particular the prediction of subcellular location, which the authors suggest might be due to some of the original sequence derived functional information being lost, despite precautions to prevent this. Nonetheless, ProstT5 is proposed as one of the early attempts toward a structure, function, and evolutionary multi-modal protein language model, with authors adding that conditional factors that specify function (e.g. GO terms) could be a potential future avenue of improvement [15].

We thus propose that combining three complementary PLMs, each pre-trained on different tasks, would provide a performance on enzyme function prediction that exceeds that of any individual encoder: ESM2 would provide information on evolutionary co-variation, ProstT5 on fold topology, and ProteinBERT, which has been jointly pre-trained on sequences and gene ontology (GO) annotation, would provide function-aware representations [17]. However, a naive concatenation of these PLMs’ embeddings would not suffice, since each PLM encodes information on a different scale and orientation. In the concatenated vector, distances would be dominated by the highest-magnitude component, and the lower-magnitude signals would be drowned out rather than provide a performance increase [18]. A possible solution to overcome this would be to normalize the individual components, resulting in them all having equal magnitude. However, this would not address their varied geometries, and more crucially, not allow each component to be weighted by functional importance. Hence, a learned projection onto the concatenated embeddings would be required to map out the combined space, such that the distance corresponded to enzyme function.

Thus, we present FuncSeek, a multi-PLM contrastive learning system for enzyme functional similarity search. FuncSeek combines ESM2, ProstT5, and ProteinBERT embeddings into a single 2816-dimensional input and projects it through a supervised contrastive head to a 256-dimensional search space optimized for EC-number retrieval. We demonstrate substantial improvements over CLEAN, BLASTp, and Foldseek on the CARE sets for remote homologs and multi-functional enzymes, while also finding that consistent misclassification is a remaining challenge for all methods we evaluate. We then show that the learned projection can be applied without further fine-tuning to a TrEMBL-scale database and that FAISS (Facebook AI Similarity Search) nearest-neighbor search allows for millisecond-scale queries on unreviewed protein collections.

## Results

### FuncSeek outperforms CLEAN on the CARE benchmark

We evaluated FuncSeek on the CARE sets (Fig 1, Table 1). CLEAN and Foldseek baselines were taken from Table 2 of Yang et al. [14]. We independently reproduced the Diamond BLASTp protocol and obtained identical results to those reported by Yang et al. [14]. Each set tests a specific failure mode: distant homologies (*ood30*), modest homologies (*ood30_50*), activity cliffs (*Price*), and multifunctional enzymes (*promiscuous*). We assess FuncSeek with both centroid and nearest-neighbour (NN) retrieval.

**Fig 1.**
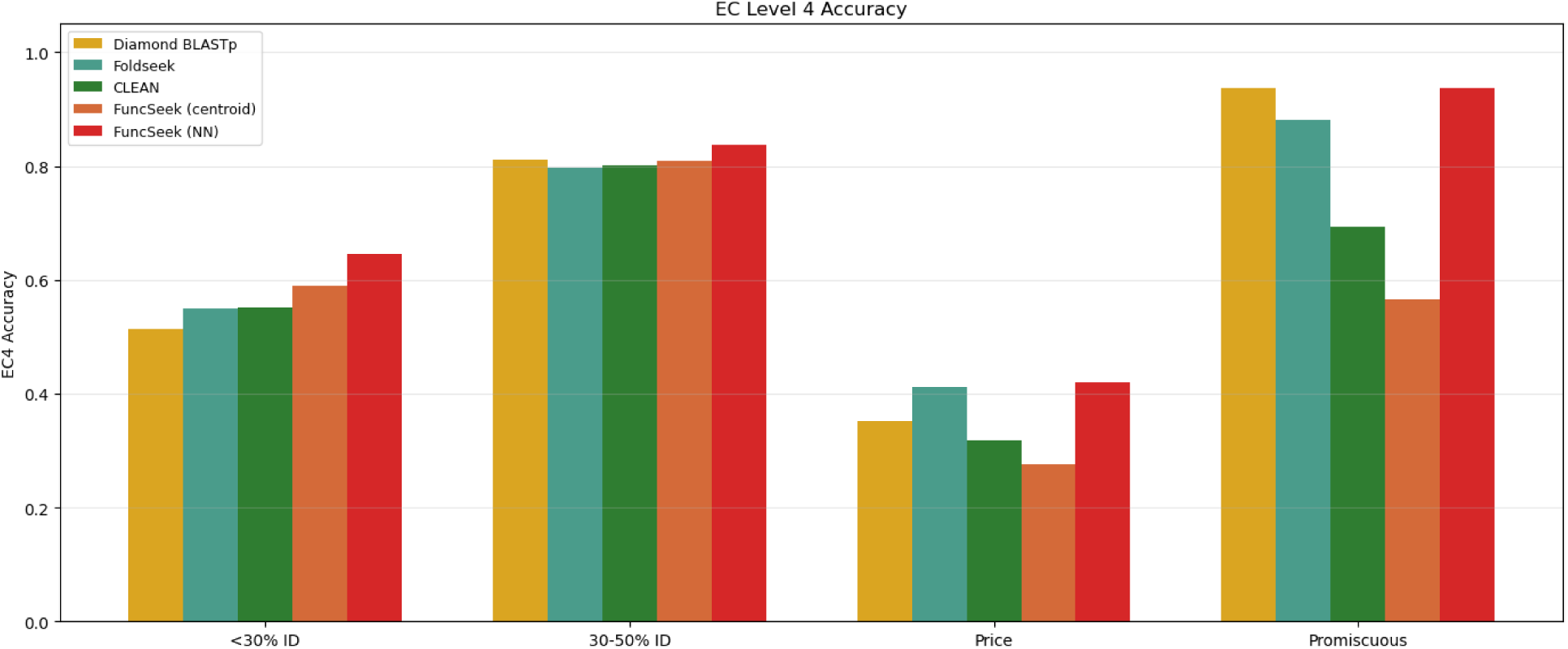
Top-1 EC4 accuracy on the four CARE benchmark sets. Diamond BLASTp, CLEAN and Foldseek values are taken from Table 2 of Yang et al. [14]. FuncSeek results are reported for centroid retrieval (using the same inference protocol as CLEAN) and for retrieval of the nearest-neighbour (NN) within all training proteins.

**Table 1.** Top-1 EC4 accuracy (%) on CARE benchmark sets.

|  | **ood30** | **ood30_50** | **Price** | **promiscuous** |
| --- | --- | --- | --- | --- |
| Diamond BLASTp | 51.4 | 81.1 | 35.1 | **93.7** |
| Foldseek | 54.9 | 79.8 | 41.2 | 88.0 |
| CLEAN | 55.1 | 80.2 | 31.8 | 69.4 |
| *FuncSeek (centroid)* | *59.0* | *80.9* | *27.7* | *56.5* |
| *FuncSeek (NN)* | **64.6** | **83.8** | **41.9** | **93.7** |

**Table 2.** Precision@K (EC4) on CARE validation set (18,293 queries vs 153,888 training index).

|  | **P@1** | **P@10** | **P@100** | **P@1000** |
| --- | --- | --- | --- | --- |
| ESM2 | 98.9 | 93.7 | 80.3 | 51.6 |
| ProstT5 | 98.4 | 95.8 | 83.8 | 61.6 |
| ProteinBERT | 97.6 | 94.0 | 86.1 | 66.6 |
| Concat (unnorm) | 99.0 | 94.6 | 82.4 | 55.8 |
| Concat (norm) | 99.0 | 94.6 | 82.4 | 55.8 |
| Per-model norm | 98.7 | 95.0 | 86.7 | 67.9 |
| CLEAN | 99.4 | 96.0 | 88.9 | 69.0 |
| *FuncSeek* | **99.5** | **97.9** | **96.4** | **96.5** |

Overall, FuncSeek had a nearest-neighbour accuracy (fraction of queries whose top-1 prediction matches the true EC4 label) of 64.6% on the *ood30* set (432 proteins, <30% identity to any training protein), while CLEAN, Foldseek and Diamond BLASTp perform at 55.1%, 54.9% and 51.4% respectively. Even when restricted to the centroid-only approach (as CLEAN uses), FuncSeek still scores 59.0%, exceeding CLEAN’s 55.1% by 3.9 percentage points. The nearest-neighbour performance for FuncSeek on the *ood30_50* set was 83.8%, compared to 80.2% for CLEAN, 81.1% for BLASTp and 79.8% for FoldSeek.

On the promiscuous set (209 multi-functional enzymes), FuncSeek nearest-neighbour retrieval achieved 93.7% EC4 accuracy, equaling Diamond BLASTp (93.7%) and significantly outperforming CLEAN (69.4%). This is mainly because centroid-based retrieval is not suitable for multi-functional enzymes, as it returns only one EC4 classification, whereas these proteins carry multiple EC annotations. As such, FuncSeek centroid retrieval obtained only 56.5% EC4 accuracy. However, nearest-neighbour retrieval handles multi-label prediction intrinsically, inheriting the entire annotation set of the closest neighbour, equivalent to the approach BLASTp uses [5, 19], although in learned embedding space rather than sequence space.

On the *Price* set (148 proteins near activity cliffs), FuncSeek nearest-neighbour retrieval had 41.9% accuracy, which outperformed CLEAN (31.8%) and Foldseek (41.2%). Centroid retrieval performed poorly overall (FuncSeek centroid: 27.7%) as a centroid is the mean over all members of a class, including legacy misannotations, which could shift it away from the true functional cluster. Nearest-neighbour retrieval is more tolerant of this effect, as long as at least one correctly annotated neighbor exists.

### Multi-PLM fusion outperforms individual models and naive concatenation

To assess the model contribution of each embedding, we tested the eight input representations (ESM2, ProstT5, ProteinBERT, unnormalized concatenation, L2-normalized concatenation, per-model normalized concatenation, CLEAN, and FuncSeek) on the CARE validation set (18,293 withheld proteins against the 153,888 training index). This allows us to isolate the contribution of each PLM and determine whether the learned fusion extracts complementary signal beyond what any single encoder provides. We compared Precision@K (P@K) at the four-digit, fourth-level EC number (EC4) (proportion of K best-ranked neighbors that have the same EC4 label as the query). A query is eligible only if its EC4 class has at least K members in the index. All representations are evaluated using the same nearest-neighbour retrieval protocol, including the CLEAN projection, to isolate embedding quality from inference strategy.

In terms of individual PLM ranks (see Table 2 and Fig 2): ProstT5 was best at 95.8% (P@10), followed by ProteinBERT (94.0%) and ESM2 (93.7%). At larger K values, ProteinBERT (86.1% P@100, 66.6% P@1000) surpasses ProstT5 (83.8%, 61.6%), while ESM2 continues last (80.3%, 51.6%). ProteinBERT’s strong performance at large K suggests that GO-aware pretraining produces functionally coherent neighbourhoods that persist deep into the ranking, though its lower dimensionality (512-d vs 1024-d/1280-d) may also contribute to denser clustering.

**Fig 2.**
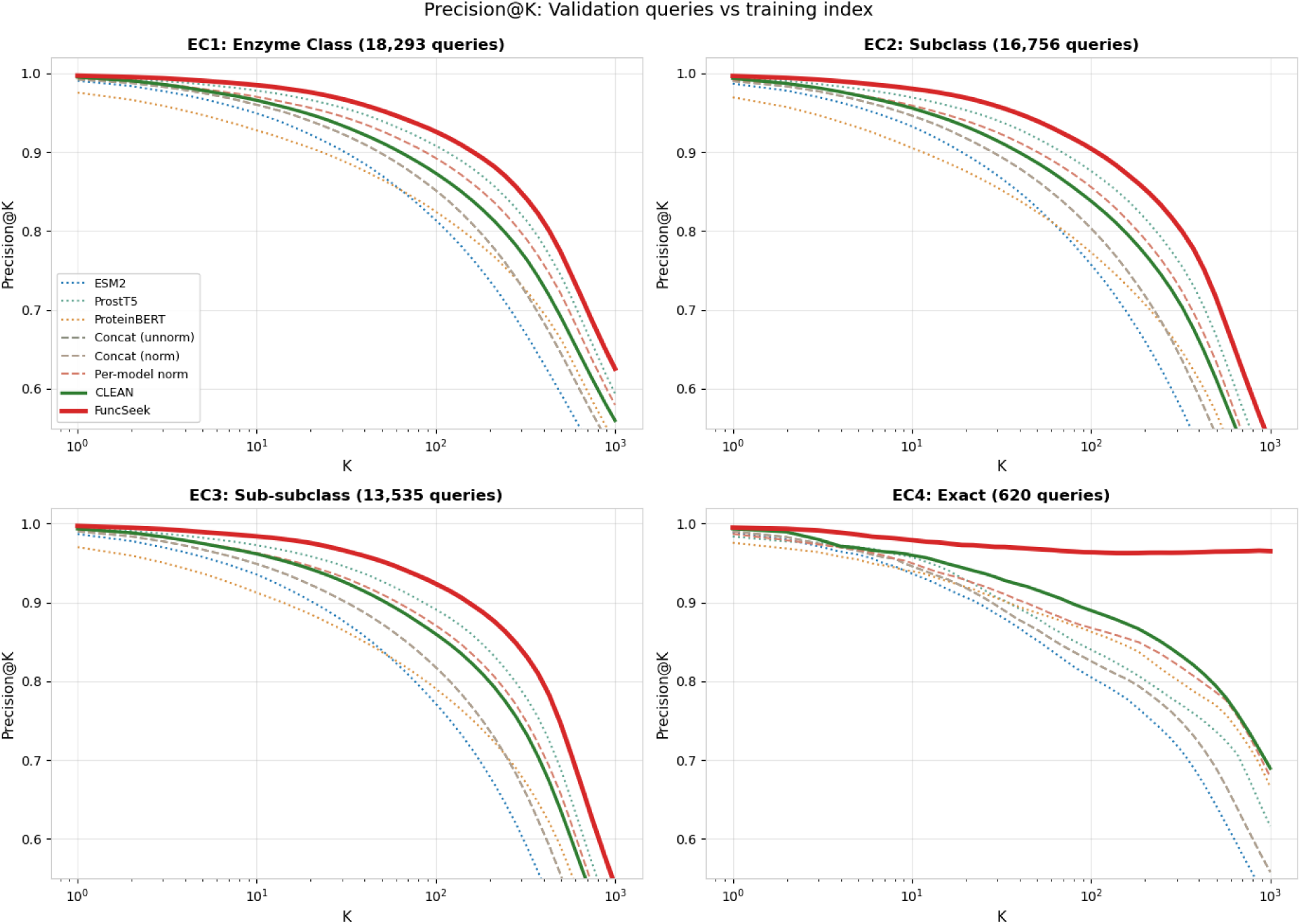
Precision@K for validation queries (18,293 proteins) searched against training index (153,888 proteins) at levels 1-4 of the EC hierarchy. For a given K, a query is eligible if its EC class has at least K members in the index. FuncSeek retains almost flat precision at depth (96.5% at EC4, K=1000) where all baselines deteriorate beyond K=100.

The concatenation of raw PLM embeddings (unnormalized or L2-normalized) achieves identical 94.6% (P@10), since cosine retrieval is scale-invariant, making global normalization redundant. Both are better than ESM2 alone (93.7%) but worse than ProstT5 (95.8%). At K=1000, simple concatenation (55.8%) falls below ProstT5 (61.6%) and ProteinBERT (66.6%), suggesting that concatenating different models’ raw representations introduces dimensional interference that degrades retrieval at depth. Normalizing each model before concatenation enhances the performance of concatenated models (67.9% P@1000) though it still loses quality compared to CLEAN (69.0% P@1000).

Pairwise ablation further demonstrates this dimensional interference (Table 3, Fig 3). Concatenation of any two of the PLMs with no learned fusion performs worse than the best individual model for Precision@1000, where ESM2 + ProstT5 (55.7) is worse than ProstT5 (61.6) alone, ESM2 + ProteinBERT (51.9) is worse than ProteinBERT (66.6) alone, ProstT5 + ProteinBERT (62.1) is worse than ProteinBERT (66.6) alone. Even all three signals together (55.8) does not surpass the best of the pair combinations. Only the learned projection FuncSeek leverages all three.

**Fig 3.**
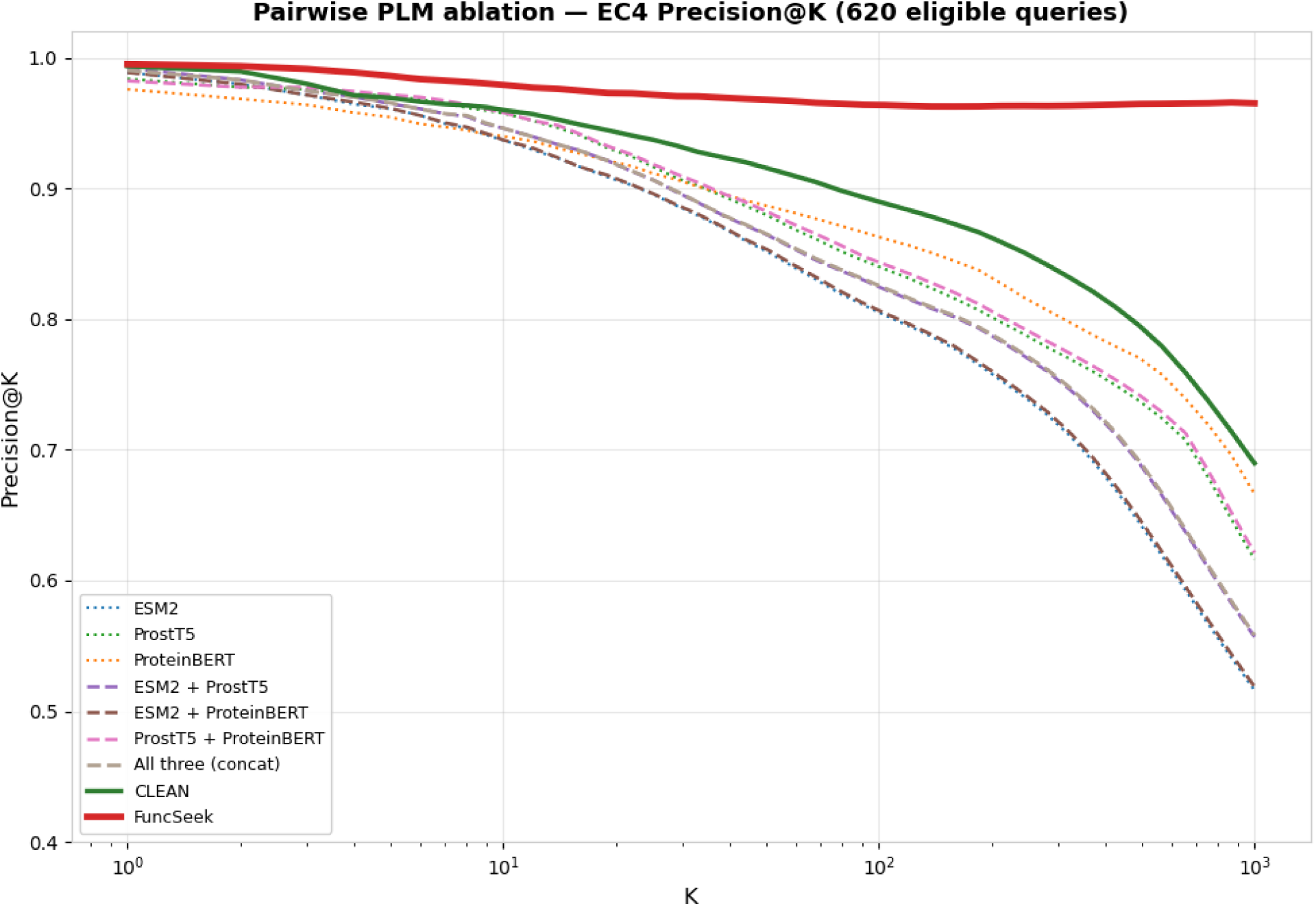
Pairwise PLM ablation. EC4 Precision@K for individual PLMs, all pairwise concatenations, the full three-way concatenation, CLEAN, and FuncSeek. Pairwise concatenations (dashed) consistently underperform the best individual model in each pair at large K, confirming dimensional interference from naively combining heterogeneous embedding spaces. FuncSeek (red, solid) is the only method that successfully fuses all three signals.

**Table 3.** Single-PLM contrastive ablation - EC4 Precision@K (620 eligible validation queries vs training index).

|  | P@1 | P@10 | P@1000 |
| --- | --- | --- | --- |
| ESM2 (raw) | 98.9 | 93.7 | 51.6 |
| ESM2 (contrastive) | 99.2 | 96.8 | 87.1 |
| ProstT5 (raw) | 98.4 | 95.8 | 61.6 |
| ProstT5 (contrastive) | 99.2 | 96.5 | 81.8 |
| ProteinBERT (raw) | 97.6 | 94.0 | 66.6 |
| ProteinBERT (contrastive) | 97.7 | 93.6 | 66.6 |
| CLEAN (ESM-1b, contrastive) | 99.4 | 96.0 | 69.0 |
| *FuncSeek (all 3, contrastive)* | **99.5** | **97.9** | **96.5** |

The trained FuncSeek projection attained 97.9% (P@10) and 96.5% (P@1000), respectively, surpassing all baselines. FuncSeek and CLEAN are almost identical at K=1 (99.5% vs 99.4%), while they separate drastically as K increases: 7.5 points apart at P@100 (96.4% vs 88.9%), and 27.5 points apart at P@1000 (96.5% vs 69.0%). While the precision of CLEAN gradually decreases after K=100, FuncSeek maintained precision (Fig 2), confirming that the multi-PLM projection creates more concentrated EC4 clusters that maintain their integrity at depth.

### Contrastive training on a single PLM does not reach FuncSeek performance

To determine whether the performance of FuncSeek is attributable to the integration of multiple PLMs, or solely due to the process of contrastive training (i.e. the learned projection from raw embeddings into the retrieval space), we trained the same projection head on frozen ESM2 (1280-dimensional input), ProstT5 (1024-dimensional input) and ProteinBERT (512-dimensional input) embeddings, respectively, using the same training hyperparameters, loss function and sampling strategy (Table 4, Fig 4).

**Fig 4.**
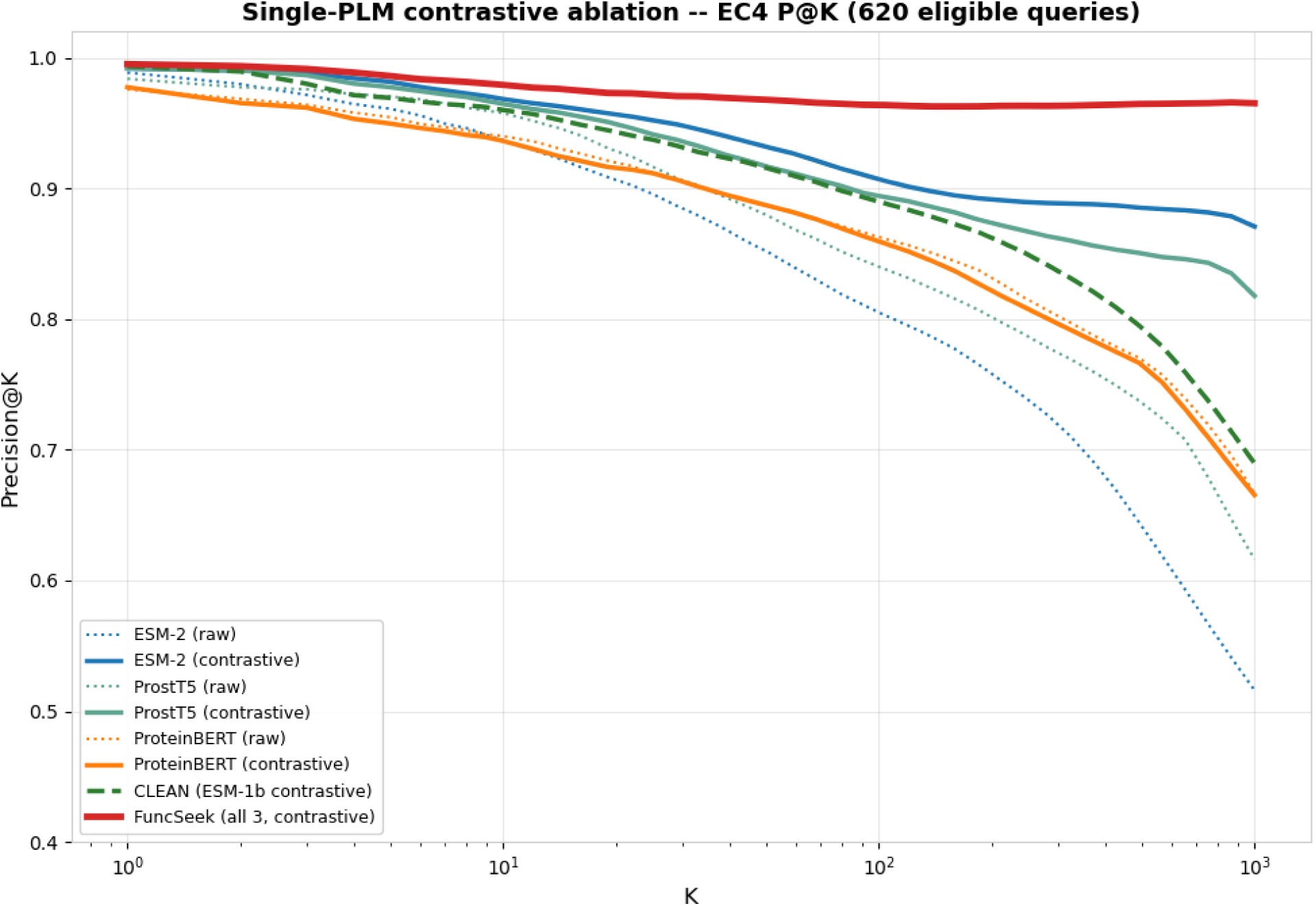
Single-PLM contrastive ablation. EC4 Precision@K comparing raw PLM embeddings (dotted lines) with PLMs trained with contrastive learning using the same architecture and hyperparameters (solid line, same color). ESM2 alone, trained with contrastive learning, achieves a 35-point boost over raw embeddings at K=1000 (51.6% to 87.1%) and significantly improves over CLEAN (69.0%), although the performance is still behind that of FuncSeek (96.5%), which indicates there is additional signal from multiple PLMs.

**Table 4.** Pairwise PLM ablation - EC4 Precision@K (620 eligible validation queries vs training index).

|  | **P@1** | **P@10** | **P@1000** |
| --- | --- | --- | --- |
| ESM2 | 98.9 | 93.7 | 51.6 |
| ProstT5 | 98.4 | 95.8 | 61.6 |
| ProteinBERT | 97.6 | 94.0 | 66.6 |
| ESM2 + ProstT5 | 99.2 | 94.6 | 55.7 |
| ESM2 + ProteinBERT | 98.9 | 93.7 | 51.9 |
| ProstT5 + ProteinBERT | 98.2 | 95.8 | 62.1 |
| Concat (norm) | 99.0 | 94.6 | 55.8 |
| CLEAN | 99.4 | 96.0 | 69.0 |
| *FuncSeek* | **99.5** | **97.9** | **96.5** |

Contrastive learning greatly improved ESM2 and ProstT5 performance, with a Precision@1000 increase of 35.5 and 20.2 percentage points, respectively. This suggests that supervised contrastive learning can indeed re-arrange the embedding space appropriately for functional retrieval. Nevertheless, the best single PLM contrastive model ESM2 (87.1%) was still 9.4 percentage points below FuncSeek (96.5%), demonstrating that multi-PLM fusion could provide benefits beyond single-PLM contrastive training. ProteinBERT exhibits no improvement from contrastive training (66.6%), possibly because 512 dimensions encode less discriminative information to begin with.

### Val-vs-val retrieval confirms generalisation beyond training memorisation

To confirm that the reported performance does not simply stem from memorisation of the training set layout, we constructed an index using solely the 18,293 validation proteins and executed nearest-neighbour retrieval on it (omitting the query protein itself). All of the validation proteins were withheld from training, and did not contribute towards any parameter updates.

FuncSeek outperformed all baselines at all values of K (Table 5, Fig 5), indicating that the learned metric can generalize to unseen proteins. Precision values are generally lower than when querying the full training index, because of the reduced index size (and thus, fewer available same-EC candidates to retrieve). Naive concatenation (38.6% at P@50) is worse than both ProteinBERT (40.6%) and ProstT5 (47.8%), again confirming dimensional interference. FuncSeek (54.6%) exceeded the best individual PLM (ProstT5, 47.8%) by 6.8 percentage points, demonstrating that the learned fusion provides genuine complementary signal beyond any single model.

**Fig 5.**
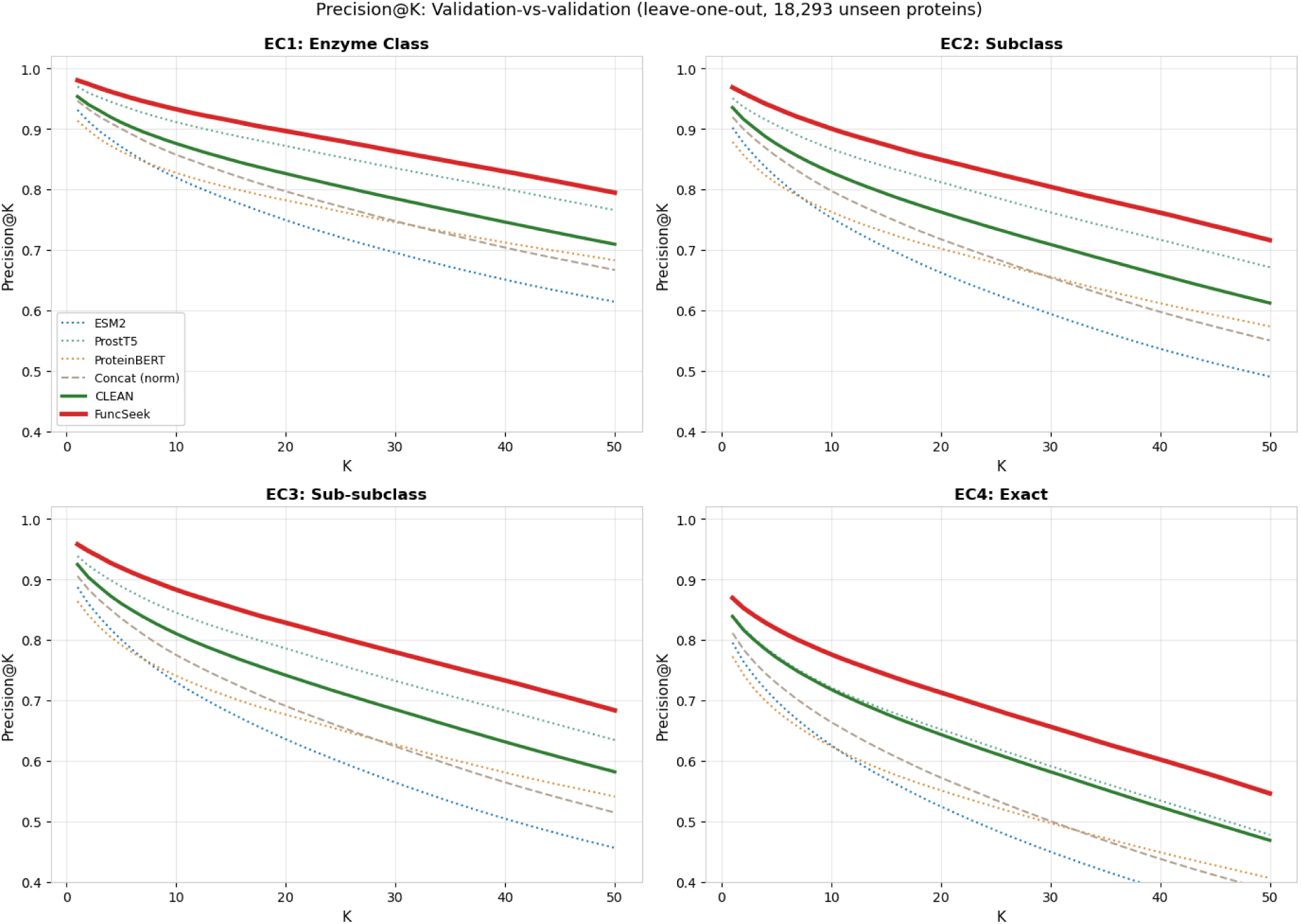
Precision@K for validation-only set (self-excluded retrieval, 18,293 unseen proteins). The index and queries are taken from the validation set only (no proteins are from the training set). FuncSeek obtained higher values for all K, which suggests the learned metric doesn’t simply memorise the training set, but generalises to unseen proteins. The lower absolute precision compared to Fig 2 is due to the smaller index size.

**Table 5.** Val-vs-val Precision@K (EC4 level, self-excluded retrieval, 18,293 unseen proteins).

|  | P@1 | P@50 |
| --- | --- | --- |
| ESM2 | 79.5 | 33.9 |
| ProstT5 | 84.0 | 47.8 |
| ProteinBERT | 77.2 | 40.6 |
| Concat (norm) | 81.1 | 38.6 |
| CLEAN | 83.9 | 46.9 |
| **FuncSeek** | **86.9** | **54.6** |

### Learned representations transfer to TrEMBL without retraining

In order to assess generalisation to the broader protein universe, we also measured the performance of FuncSeek on a dataset of 8,031 TrEMBL entries that could be mapped to BRENDA. This data set represents enzymes that have been experimentally studied and for which kinetic data have been published; hence this represents a highly confident set of annotations that does not solely rely on computational transfer of annotations. None of these were encountered in the training set. We performed nearest-neighbour retrieval (self-excluding) where a query is eligible only if at least K other proteins sharing its EC annotation exist in the index; at K=10 this yields 181 EC4-level eligible classes and 4,269 eligible queries (Table 6, Fig 6):

**Fig 6.**
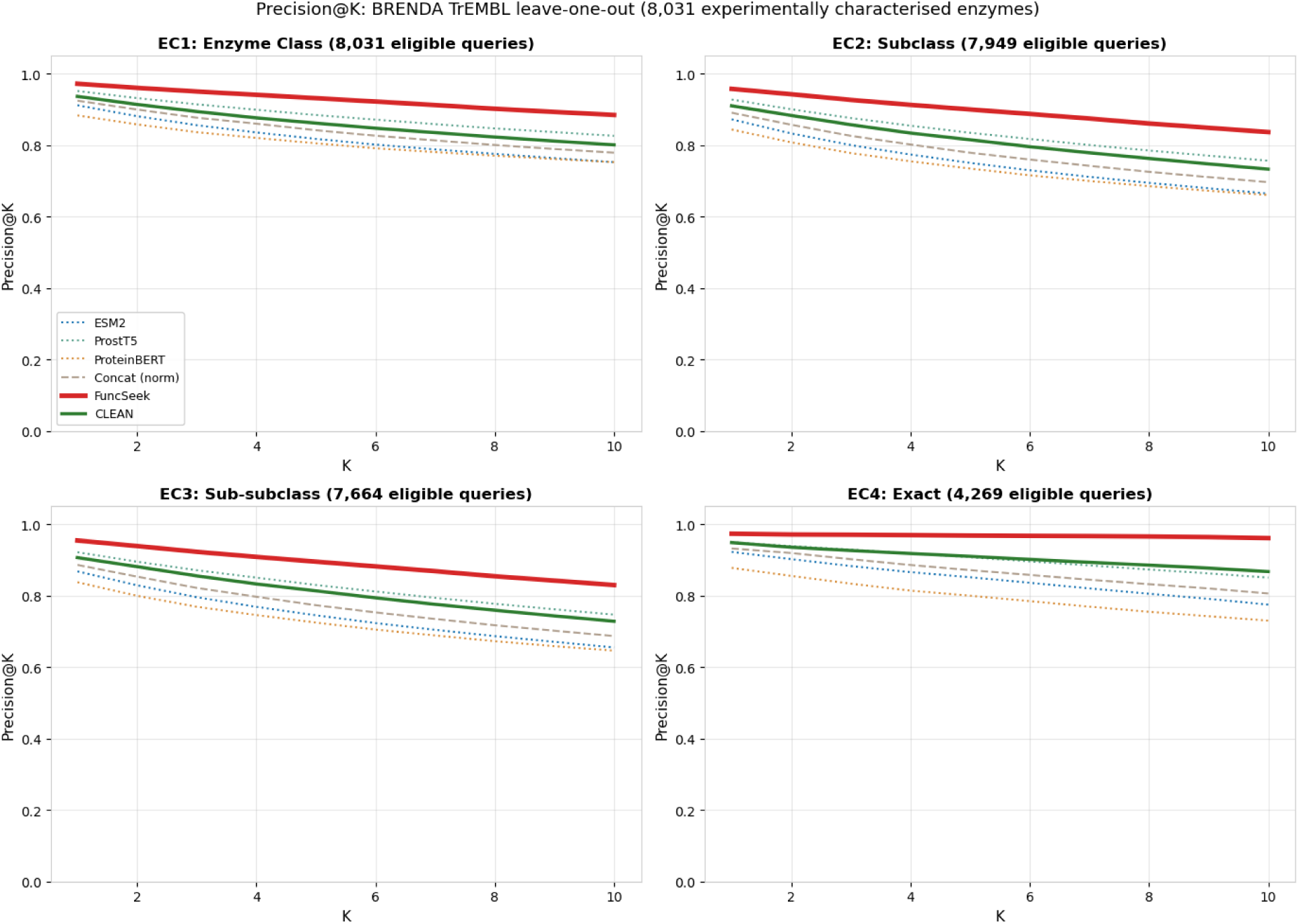
Precision@K for BRENDA-validated TrEMBL proteins (8,031 characterized proteins; self-excluded retrieval, no overlap between training and testing sets). FuncSeek maintains the highest precision across all EC levels and values of K; the learned projection generalizes to proteins not present in the SwissProt training set without further training.

**Table 6.** BRENDA TrEMBL Precision@K (EC4 level, self-excluded retrieval).

|  | P@1 | P@5 | P@10 |
| --- | --- | --- | --- |
| *FuncSeek* | **97.3** | **96.8** | **96.1** |
| CLEAN | 94.8 | 91.0 | 86.7 |
| ProstT5 (raw) | 94.9 | 90.7 | 85.0 |
| ESM2 (raw) | 92.2 | 85.1 | 77.5 |
| concat_norm | 93.2 | 87.1 | 80.6 |
| ProteinBERT | 87.8 | 80.0 | 73.0 |

FuncSeek outperforms all baselines at all values of K, and the gap widens at higher K: at Precision@10, FuncSeek exceeds CLEAN by 9.4 percentage points (96.1% vs 86.7%). This suggests the learned projection does not just position the single nearest-neighbor correctly, but is able to arrange entire EC4 classes as compact clusters in the embedding space. The precision is maintained throughout the ranked list, even on unseen data.

Disaggregation at Precision@1 showed asymmetrical errors (only 70 instances of CLEAN being correct where FuncSeek is not, vs 482 in the opposite direction). Out of the 70 errors made by FuncSeek, 78.6% returned a protein in the same EC level 3 class (same reaction type, incorrect substrate specificity), qualifying them as near-misses rather than outright failures.

## Discussion

The major result presented in this paper is that three PLMs with different training tasks embed biological signals such that when mapped with a learned projection into a unified retrieval space, the fusion method outperforms any single encoder. On the *ood30* set (the most challenging, remote homology), FuncSeek nearest-neighbour retrieval (64.6%) performs 9.5 percentage points better than CLEAN’s centroid prediction (55.1%), and 13.2 percentage points better than Diamond BLASTp (51.4%). FuncSeek also matches BLASTp on promiscuous enzymes (93.7%), whereas CLEAN’s centroid retrieval is at 69.4%. Two possible explanations are proposed for FuncSeek’s performance: (1) The combination of PLMs with a learned projection can lead to a more expressive embedding space than a single encoder. (2) Nearest-neighbour search can better transfer annotation information than centroid-matching, especially where enzymes have multiple EC annotations or class centroids are corrupted by false annotations. FuncSeek’s own centroid retrieval (59.0% on *ood30*) also outperforms CLEAN, but nearest-neighbour retrieval adds a further 5.6 percentage points, confirming that retrieval strategy contributes independently of embedding quality.

### Why multi-PLM fusion succeeds

Raw concatenation of ESM2, ProstT5 and ProteinBERT results in a 2816-dimensional vector that performs worse than the best individual PLMs. This is likely because each PLM embeds proteins in a different geometric space with different scale and metric, and when comparing proteins using the raw concatenation the distances are defined by the strongest signal (i.e. the embedding with highest L2 norm) instead of by the signal which carries the most information. Pairwise ablation provides direct evidence of this dimensional interference. For each of the pairwise concatenations, performance is worse than the better of the two models paired up (at K=1000): ESM2 + ProstT5 (55.7%) < ProstT5 (61.6%), ESM2 + ProteinBERT (51.9%) < ProteinBERT (66.6%), ProstT5 + ProteinBERT (62.1%) < ProteinBERT (66.6%). The full three-way concatenation (55.8%) performs no better than the best pairwise combination (62.1%), showing that even with increasing numbers of models in the combination, naive concatenation cannot improve retrieval accuracy at depth.

The contrastive projection head learns to rotate and reweight each PLM embedding space such that cosine similarity can be used for EC4 membership. On the CARE validation set, FuncSeek has EC4@1000 = 96.5% while CLEAN only gets 69.0% and the best raw PLM (ProteinBERT) gets 66.6%. This increased separation at large K is a hallmark of a properly learned metric space. While all methods perform well at shallow retrieval, maintaining precision deep into the ranked list requires true separation between class boundaries. This separation generalises beyond the training distribution: on BRENDA TrEMBL (8,031 experimentally characterised enzymes, zero overlap with training), FuncSeek achieves 96.1% at K=10, 9.4 points above CLEAN (86.7%).

Now, a natural objection that could arise is that contrastive learning alone causes this improvement, not multi-PLM fusion. Thus, we tested this hypothesis directly. The same projection head trained on ESM2 alone improves Precision@1000 from 51.6% to 87.1%; over 35 percentage points (Table 4). This clearly demonstrates supervised contrastive learning can dramatically transform the embedding space of any PLM. However, even the best single-PLM contrastive model (ESM2, 87.1%) is still 9.4 percentage points behind FuncSeek (96.5%). ProteinBERT does not benefit from contrastive learning, presumably because its embedding space is fundamentally more constrained (512 dimensions). Contrastive training of ESM-1b (CLEAN, 69.0%) clearly shows that the encoder is also important. The remaining 9.4-point gap can then likely be attributed to the effect of the multi-PLM fusion, where signal, which is not individually present within any single PLM, is successfully extracted by the projection function.

ProstT5’s strength among raw PLMs was somewhat surprising, as we expected the largest corpus-trained PLM, ESM2, to perform best at neighbourhood purity. The raw embedding ranking (ProstT5 61.6% vs ESM2 51.6% at P@1000) suggests that bilingual sequence-structure pretraining produces representations that are natively better organised for functional retrieval, likely because encoding 3Di states forces the model to capture active-site geometry and substrate-binding pocket topology [15]. However, contrastive training reverses this ranking: ESM2 contrastive (87.1%) surpassed ProstT5 contrastive (81.8%), indicating that the functional signal was already latent in ESM2’s representations but required a learned projection to expose it. The structural encoding thus provides a head start in raw space, not a ceiling advantage once the embedding space is explicitly optimised for function.

### Nearest-neighbour retrieval vs centroid retrieval

FuncSeek’s nearest-neighbour retrieval consistently outperforms centroid matching, both its own (FuncSeek centroid, Table 1) and CLEAN’s. CLEAN assigns a query to its nearest class centroid, where each centroid is the average embedding of all training proteins in that EC4 class [5]. This collapses all within-class structure into a single representative, making the prediction vulnerable to both misannotation among class members and within-class diversity. BLASTp avoids this by transferring all annotations from the best-hit sequence, which naturally handles multi-functionality. We apply the same nearest-neighbour logic in learned embedding space.

The FuncSeek results demonstrate this clearly. Centroid retrieval performance fails completely on the promiscuous set (56.5%) compared to nearest-neighbour retrieval performance of 93.7%. A promiscuous enzyme’s nearest neighbour in embedding space is likely another multi-functional enzyme carrying a similar set of EC annotations, so the full annotation set transfers naturally. Centroid matching requires the query to simultaneously rank near multiple independent class centroids, which is geometrically more demanding and dependent on the max-separation heuristic. Similarly, centroid retrieval performs substantially worse on the *Price* set (27.7%) compared to the nearest-neighbour retrieval (41.9%). This is likely because an EC class includes some members that have retained outdated (erroneous) annotation (e.g., a correctly reannotated protein), and the centroid may have absorbed the incorrect information. In contrast, the nearest-neighbour approach only depends on having found at least one representative with correctly retained annotation.

The choice between the paradigms is thus based on the application; centroid for rapid classification of single-function enzymes and nearest-neighbour for transfer annotation where multi-functionality and full coverage of annotation are important.

### Activity cliffs define the boundary of sequence-derived representations

The *Price* set (148 enzymes near activity cliffs, where small sequence changes alter substrate specificity) is where all methods struggle. FuncSeek nearest-neighbour retrieval (41.9%) is marginally above Foldseek (41.2%) and Diamond BLASTp (35.1%), but well below the performance achieved on other splits. Centroid retrieval fares even worse (FuncSeek centroid 27.7%, CLEAN 31.8%), consistent with the vulnerability to misannotation discussed above. The EC-level breakdown reveals where the failure concentrates: FuncSeek achieves 95.9% at EC1 and 93.2% at EC2, but drops to 41.9% at EC4. The bottleneck is likely substrate specificity and stereochemistry (the fourth EC digit), precisely the information encoded in active-site geometry. None of the input representations, whether sequence co-variance (ESM2), predicted fold topology (ProstT5), or GO-informed sequence (ProteinBERT), explicitly encode binding-pocket geometry at residue resolution. Discriminating enzymes that differ by a handful of catalytic residues likely requires explicit modelling of these interactions, which seems beyond the reach of the current PLM embeddings.

### Transfer to TrEMBL confirms generalisation

The BRENDA TrEMBL benchmark (8,031 enzymes with experimentally validated EC annotations from kinetic data, none observed during training) serves as the most convincing test of FuncSeek’s out-of-distribution generalisation. FuncSeek retains its advantage over CLEAN on these unseen, experimentally characterised enzymes, and the margin widens with retrieval depth. The error analysis is more revealing than the headline accuracy: FuncSeek’s disagreements with CLEAN are strongly asymmetric, and where FuncSeek does err, the great majority of cases return an enzyme of the correct reaction type but incorrect substrate specificity. These are near-misses at the finest level of the classification rather than broad functional failures, which points to substrate discrimination, not reaction class, as the residual bottleneck.

### Limitations

There are a number of factors that restrict our analysis and interpretation. Firstly, we are using EC labels derived from SwissProt which, although manually curated, is not immune to annotation error [4]; the model can only be as good as the data it is trained on. Secondly, by using only the primary EC annotation per protein during training, we inherently disregard multi-functionality. Although we assess multi-label prediction via nearest-neighbour retrieval (inheriting annotations from the K nearest neighbours), the training signal is that of only one EC per protein. We expect further improvements would arise from an alternative training objective that specifically addresses multi-functional proteins. Finally, the projection head is trained on embeddings from fixed PLMs. As models are retrained or improved, the embeddings may change and the projection head will need to be re-trained, although with only ∼1.8M parameters this is not a significant cost.

### Positioning and future directions

We view FuncSeek as complementary to structure-based methods. On activity cliffs (*Price* set), FuncSeek nearest-neighbour retrieval (41.9%) performs at a similar level to Foldseek (41.2%), and both outperform centroid retrieval in these cases. Where alignment methods perform poorly (distant homologs with less than 30% identity or multi-functional enzymes), FuncSeek provides large improvements that Foldseek does not. An ensemble approach that defaults to FuncSeek for general annotation but defers to structure-based methods when the nearest neighbours span multiple EC4 classes could retain the advantages of each.

The general principle is that multi-PLM fusion by contrastive projection is applicable to any protein property for which labelled training data exists. The pre-computed PLM embeddings are independent of the downstream task, so generalisation from EC numbers to Gene Ontology categories, substrate specificity classes, or thermostability bins requires only a change in the positive-pair criterion of the contrastive loss; the output dimensionality (256) and retrieval infrastructure remain constant. Perhaps most importantly, the approach can annotate proteins in large unannotated databases (e.g. metagenomic datasets) since only projected embeddings need to be present in the target index, not existing functional annotations. Since the projected embeddings are compact (256 dimensions) and retrieval relies on standard vector similarity search (e.g. FAISS [20]), the approach scales to billion-entry indices, making functional annotation of entire metagenomic databases feasible without per-query alignment.

## Materials and Methods

### Embedding generation

FuncSeek works by utilizing fixed, precomputed embeddings derived from three PLMs, each of which was trained with a different objective, and therefore encodes a unique aspect of protein biology. No PLM parameters are fine-tuned, and each acts as a fixed feature extractor only.

Trained on UniRef50 sequences using masked language modelling, ESM2 (facebook/esm2_t33_650M_UR50D) creates 1280-dimensional embeddings encoding evolutionary co-variation [11]. Each protein sequence is tokenized with at most 1024 amino acid residues (longer sequences are truncated). Hidden states are then obtained from the last layer and mean pooled over residue positions to produce the final representations.

ProstT5 (Rostlab/ProstT5) is a bilingual encoder-decoder that can translate between amino acid sequences and Foldseek’s 3Di structural alphabet [16]. Its output is a 1024 dimensional embedding representing three-dimensional fold topology. To generate the required representations, input sequences are prefixed with the <AA2fold> tag, separated by spaces between residues, and the encoder output mean-pooled across residue positions (sequences exceeding 2000 residues are truncated).

The ProteinBERT model generates a 512-dimensional embedding, which represents functional annotation learned by co-pretraining protein sequences and GO terms [17]. To produce these embeddings, we set the maximum sequence length to 2000 residues, and extract the output of the last global representation layer as the representation of the protein.

These three outputs are then concatenated without normalization, producing a 2816-dimensional (1280 + 1024 + 512) input vector for each protein. The reasoning behind not normalizing the individual PLM outputs before concatenation is that a projection head will readily handle differing relative scale and spatial orientation of each of the individual embedding spaces [21].

### Training data

The training data is drawn from the CARE benchmark [14]. This benchmark contains 184,529 protein-EC pairs corresponding to 173,550 distinct SwissProt proteins across 4,656 unique EC4 classes. Where more than one EC annotation is available for a protein, only the first is used as a label in our training data. EC classes are randomly split into training and validation sets (stratified by the EC4 class) in a 90:10 ratio. The validation and training sets contain no EC4 classes with fewer than two proteins, which would make positive protein pairs impossible to generate. Any proteins classified as test data in the CARE benchmark are completely removed from consideration, including the reference index, training and validation.

### Model architecture

The projection head consists of a 3-layer feedforward network that maps the 2816-dimensional concatenated input into a 256-dimensional retrieval space (Fig 7). LayerNorm is inserted before each activation for gradient stabilization between inputs of varied scale. L2 normalization of the output maps all the embeddings into a unit hypersphere in 256 dimensions. As a result, the distance between two embeddings for retrieval purpose is measured via cosine similarity on the unit hypersphere (which is the dot product). The model has ∼1.8 million trainable parameters.

**Fig 7.**
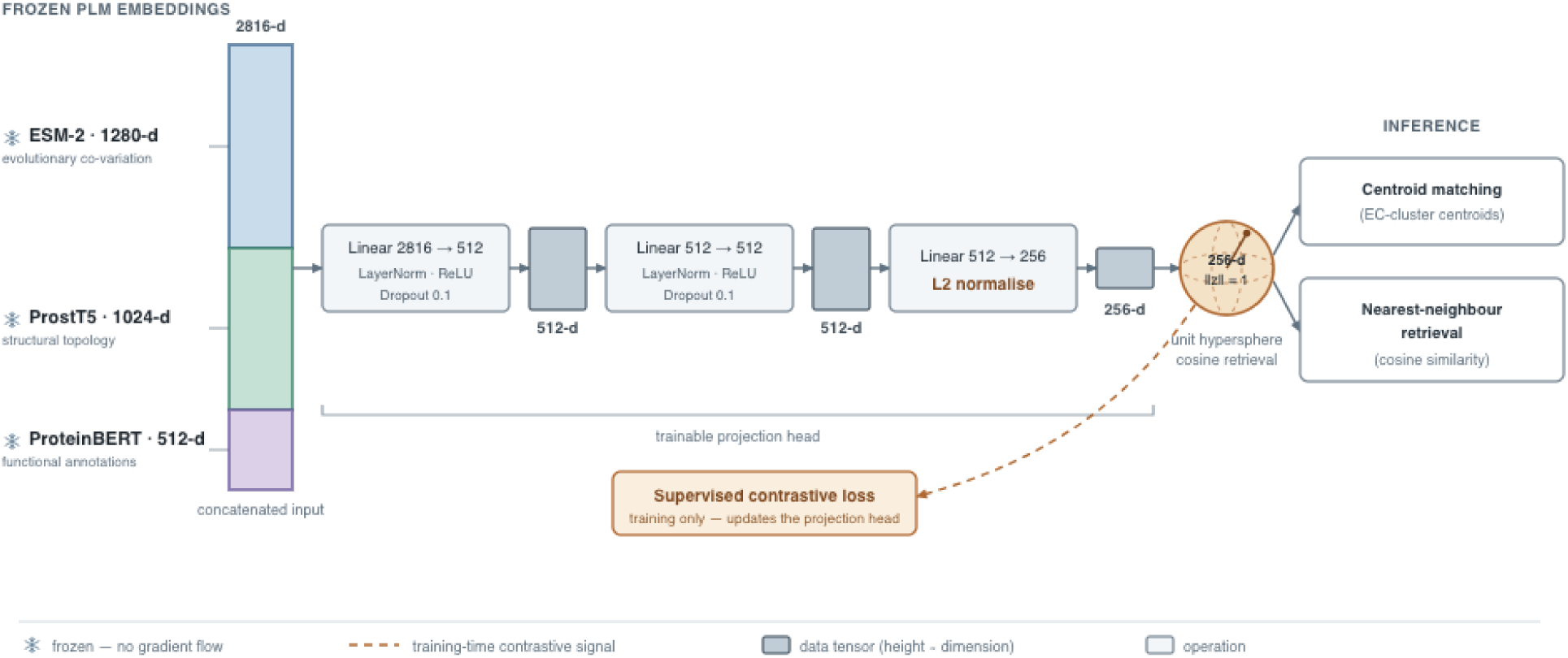
FuncSeek architecture. Frozen embeddings from ESM-2 (1280-d), ProstT5 (1024-d), and ProteinBERT (512-d) are concatenated into a 2816-dimensional input vector and passed through a 3-layer projection head that yields a 256-dimensional L2-normalized embedding on the unit hypersphere. The projection head is trained via supervised contrastive loss, and during inference the projected embedding is either used for centroid matching or K-nearest-neighbour matching based on cosine similarity.

### Loss function

To train the model, we employ supervised contrastive loss with hard negative mining (SupconH) [5] following the framework proposed by Khosla et al. [22]. Given a batch of N samples with L2-normalized projections *{z1, …, zN}* and corresponding EC4 labels *{y1, …, yN}*, the loss is given by (equation 2 in Khosla et al. [22]):

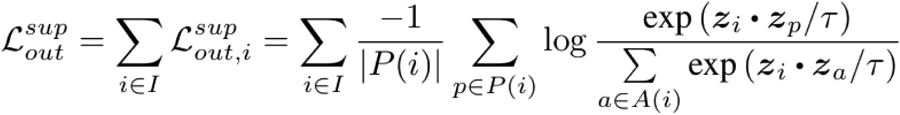

where *P(i) = {p : y p = y i, p i}* is the set of positives for anchor *i*, *A(i) = I \ {i}* is all samples except anchor *i*, and *τ = 0.1* is the temperature parameter. The loss maximizes the log-probability of each positive pair relative to all other samples in the batch, such that same-EC4 proteins become closer and different-EC4 proteins get pushed apart. Self-similarity is excluded by masking the diagonal, so that each anchor is only compared against other samples in the batch. The final loss is averaged over all positive anchors that have at least one positive in the batch.

### Batch sampling and hard negative mining

Instead of sampling randomly and uniformly, each batch is constructed in a manner intended to yield a highly informative gradient signal through a sampling strategy inspired by the authors of CLEAN [5]. Each batch consists of 64 EC4 classes; from each class, 8 positive samples are sampled (with replacement if class size is less than 8) and 30 hard negatives are sampled from the 10 nearest EC4 classes in the projected embedding space (using cosine distance between the projected embeddings). The overall effective batch size thus corresponds to *64 x (8+30)* = 2,432 samples.

Hard negatives are selected by computing a centroid for each EC4 class (the L2-normalized mean of all its members’ projected embeddings). For each class in the batch, the 10 nearest EC4 centroids by cosine distance define the hard negative pool for that class. This pool is then sampled with a probability inversely proportional to its centroid distance (closer = more likely), and a protein is drawn randomly. This process is then repeated 30 times per class, yielding 30 hard negatives that are maximally confusing for the model.

Centroids are refreshed every 50 epochs by projecting all of the training proteins through the current model, then repeating the hard negative mining described above. This dynamic updating allows “hard” to shift over the course of training: at early times, this involves protein-embedding overlap among otherwise distinct classes; at later stages, the notion of “hard” consists of EC classes that continue to cluster close to the current model’s projections.

All of the parameters relating to batch sampling (EC4 classes per batch, positives and negatives per class, and hard negative pool size) were chosen by performing Bayesian hyperparameter optimization over the search space using the Optuna hyperparameter optimization framework [23].

### Data augmentation

Input embeddings were augmented by adding Gaussian noise (mean = 0, standard deviation = 0.01) during training time (jitter augmentation). Jittering the inputs penalizes the projection head for sensitivity to minor fluctuations in PLM outputs and helps it generalize to sequences whose embeddings are somewhat outside the training distribution.

### Optimization

The Adam optimizer (learning rate = *5 × 10^−4^*, weight decay = *1 × 10^−5^*) was used to train the model. The learning rate has a cosine annealing schedule which decreases to zero by the final epoch of training (maximum 500 epochs). Training is terminated early if the validation loss does not improve for 20 epochs, keeping the checkpoint that performs best in validation. Convergence typically occurs within 50-100 epochs on a single GPU.

### Hyperparameter optimization

All architectural and training hyperparameters were determined via Bayesian optimization using Optuna (100 trials, Tree-structured Parzen Estimator sampler, and a MedianPruner to terminate unpromising runs early). A total of 11 hyperparameters encompassing model capacity, batch composition, regularization and learning rate schedule were searched. The resulting optimal configuration (hidden_dim=512, temperature=0.1, batch_n_ec=64, n_pos=8, n_neg=30, n_hard_ec=10, dropout=0.1, lr=5e-4, weight_decay=1e-5, jitter_std=0.01, centroid_refresh=50 epochs) was used for all further experiments.

### Inference

#### Centroid-based retrieval

Following the inference protocol described by Yu et al. [5], the centroid for each EC4 class is calculated as the L2-normalized mean of the projected training embeddings of that EC4 class. Then, given a query protein, its concatenated 2816 dimensional embedding is passed through the trained projection head to obtain a 256 dimensional unit vector. The cosine similarity between this vector and each of the calculated centroids is then computed and the EC4 number corresponding to the most similar centroid is taken as the function of the protein. In order to predict the function of a multi-label protein, we applied max-separation as implemented by Yu et al. [5]. This method ranks centroids by decreasing similarity and applies a cut-off at the point where the greatest difference between consecutive scores is observed, returning all EC classes above the cut-off as predictions.

#### Embedding nearest-neighbour retrieval

The cosine similarity of the query’s projected embedding to each individual training proteins’ projected embedding is measured. The EC annotations of the 10 nearest-neighbours are tallied. The fraction sharing each distinct EC number serves as a confidence score, and the most frequent EC is the predicted function. In the case of multi-functional enzymes this naturally provides multi-label prediction when the nearest-neighbour carries multiple EC annotations, analogous to homology-based annotation transfer [19]. We calculate top-1 accuracy for comparison with existing methods published in the literature [14].

### Benchmark evaluation

We test our method on the CARE benchmark [14] which defines 4 test sets under known distribution shifts. The set compositions are as follows: the 432 enzyme out-of-distribution test set (*ood30*) consists of entries with <30% sequence identity to any training protein (remote homology). The *ood30_50* test set contains 560 enzymes with 30-50% sequence identity (moderate homology), with the *Price* test set having 148 experimentally characterized enzymes that were previously incorrectly annotated in public databases. These enzymes are often located near activity cliffs, where small sequence differences could be associated with large changes in catalytic ability. The final test set consists of 209 enzymes with more than one validated EC4 annotation, which make up the *promiscuous* test set.

The projection head was trained on proteins from the CARE benchmark [14], after excluding all CARE test proteins from both the training set and reference index. The remaining proteins were split into training (153,888) and validation (18,293) sets, stratified by EC4 class, with hyperparameters identified via Optuna. We subsequently test FuncSeek with centroid- and nearest-neighbor retrieval on the embedding space, against known baselines from CLEAN, Diamond BLASTp, and Foldseek (see Yang et al. [14], Table 2).

### Component contribution analysis

To independently assess the retrieval performance of each PLM, we calculated the EC4 level Precision@K (P@K) over 8 different input representations. These include ESM2 (1280-d), ProstT5 (1024-d), ProteinBERT (512-d), simple concatenation (2816-d), concatenation with global L2 normalization, concatenation with per-model L2 normalization, the CLEAN projection and the trained FuncSeek projection (256-d). Validation proteins were used as queries against the training index (the two sets do not overlap). A query is eligible only if its EC4 class has at least K members in the index.

To further quantify each PLM’s unique signal, we also tested pairwise concatenations (ESM2+ProstT5, ESM2+ProteinBERT, ProstT5+ProteinBERT) with each combined vector L2-normalized before retrieval. If the pairwise combination is worse than the best model of the pair, this confirms that naïvely combining diverse embedding spaces introduces dimensional interference.

### Validation-set generalisation

To ascertain that retrieval quality does not arise from memorising the training set layout, we constructed an index from only the 18,293 validation proteins and performed self-excluded nearest-neighbour retrieval within this set. These proteins were completely unused during training. The same EC4 Precision@K metric is reported. The reduced index size results in generally lower absolute precision compared to the val-vs-train evaluation.

### TrEMBL generalisation: BRENDA-validated enzymes

To further evaluate if the learned projection is transferable to other proteins beyond the SwissProt training set, we also test it on a set of 8,031 TrEMBL proteins with cross-references to BRENDA [24]. These cross-references indicate experimental characterisation in the primary literature, providing high-confidence EC annotations independent of computational annotation and/or transfer. We additionally confirmed that none of these 8,031 proteins were used during training.

Each of the 8,031 proteins is searched against each of the other 8,030 proteins and Precision@K is evaluated at the EC4 level. A query is eligible at a given K if there are at least K proteins in the index with the same EC4 class (so that 100% precision can be attained). This results in 181 EC4 classes with at least 10 other members, eligible for evaluation, corresponding to 4,269 queries. Results are compared to CLEAN, individual PLMs and the concatenated normalized PLMs.

## Data availability

All code required to reproduce the results in this manuscript, including the FuncSeek training, embedding-generation, evaluation and retrieval pipelines, is openly available at https://github.com/leendertcloete/FuncSeek under the Apache License 2.0, and is archived at Zenodo (doi: 10.5281/zenodo.22149399). The same archive contains the trained FuncSeek projection weights and the projected embeddings used for retrieval.

The data underlying every reported result are provided as Supporting Information: the CARE-derived training and validation splits with their EC annotations (S1, S2), the 8,031 BRENDA-validated TrEMBL proteins used for the transfer evaluation (S3), the Precision@K and error-disaggregation tables for that evaluation (S4, S5), the Diamond BLASTp output for the four CARE test sets (S6), the trained projection heads (S7), the projected and raw embeddings needed to re-derive the reported retrieval metrics without re-running the language models (S8), and the selected hyperparameter configuration (S9).

The full SwissProt and TrEMBL embedding matrices are not deposited because of their size. They are deterministic outputs of the accession lists in S1 to S3 passed through the deposited models, and can be regenerated with the released code. The CARE benchmark is publicly available from Yang et al. (2024), doi: 10.48550/arXiv.2406.15669. The underlying protein sequences are available from UniProt (SwissProt and TrEMBL) at https://www.uniprot.org.

## Notes

http://doi.org/10.5281/zenodo.22149399

### Competing Interest Statement

The authors have declared no competing interest.

### Summary of Updates

Reformatted for journal submission. No changes to data, results or conclusions. Adds an Author Summary and a Data Availability statement, corrects figure and table numbering and placement, and reformats the references. The code and supporting data are now publicly available at https://github.com/leendertcloete/FuncSeek, archived at Zenodo (doi: 10.5281/zenodo.22149399).

